# The old pipe gives the sweetest smoke: A phylogenetic turn for eDNA metabarcoding

**DOI:** 10.64898/2026.06.11.731524

**Authors:** Rachel Haderlé, Visotheary Ung, Jean-Luc Jung

**Affiliations:** Institut de Systématique, Évolution, Biodiversité (ISYEB), Muséum national d’Histoire naturelle, CNRS, Sorbonne Université, EPHE-PSL, Université des Antilles, Paris, France; Station Marine de Dinard du Muséum national d’Histoire naturelle, Dinard, France

## Abstract

Environmental DNA (eDNA) metabarcoding has transformed biodiversity monitoring, yet most analyses rely on taxonomic metrics that are sensitive to methodological variation and limit cross-study comparability. We propose a “phylogenetic turn” in eDNA analysis through the integration of phylogenetic diversity (PD) metrics. By incorporating evolutionary relationships, PD reduces dependence on species-level resolution, increases robustness to detection biases, and better captures the evolutionary “option value” of biodiversity.

We synthesize key PD metrics across richness, divergence, and regularity, emphasizing the use of standardized effect sizes (SES) for ecological interpretation while addressing challenges in metric selection. We apply this framework to five marine eDNA datasets (2021–2025) spanning ecologically and geographically contrasting ecosystems, from tropical to Arctic regions, and encompassing a wide gradient of anthropogenic pressure.

Across datasets, we identify consistent patterns: anthropized ecosystems exhibit high taxonomic richness but reduced phylogenetic diversity, indicating phylogenetic clustering, whereas less disturbed systems show lower richness but greater evolutionary breadth. These findings demonstrate that PD reveals ecological structure not captured by taxonomic metrics, including signatures of environmental filtering and community assembly processes.

By providing a reproducible analytical workflow based on standardized eDNA datasets, we position phylogenetic diversity as a critical bridge between eDNA data and conservation frameworks. Ultimately, eDNA-based phylogenetic approaches open new avenues for decoding global biodiversity patterns across heterogeneous ecosystems.

---

Environmental DNA (eDNA) metabarcoding encompasses a variety of sampling and analytical protocols ^1–3^. In general, the primary output is a list of taxa. How this list is subsequently used varies widely: most studies focus on taxonomic richness and beta-diversity analyses, and one can also examine the functional diversity, the identity of remarkable taxa, or their variability across environmental or experimental gradients ^4–10^. These diverse approaches may often be difficult to compare across studies or datasets, typically because of their high sensitivity to experimental variables.

Reflecting this variability, standardization and data sharing of eDNA data are still infrequently used, event through their FAIRification became crucial. eDNA data can now be published following the GBIF guide for DNA-derived occurrence data ^11^ and dedicated recommendations ^12^, ensuring compliance with existing standards ^13,14^. Haderlé et al. ^15^ is an example of a standardised marine vertebrate eDNA dataset published following the FAIR principles ^16^.

Despite such standardization efforts, from experimental protocols to resulting datasets, a well-known concern in eDNA metabarcoding remains the high variability observed among samples. This variability may be driven by heterogeneous protocols (from sampling to laboratory and bioinformatic steps; ^1,3,17^), the patchy spatiotemporal distribution of eDNA, particularly in marine environments ^15,18^, partly reflected in the number of replicates required to approach sampling saturation ^19^, and persistent limitations in taxonomic assignment ^20–22^.

Taxonomic inventories derived from eDNA-based approaches may therefore exhibit significant inconsistencies, as may measures of ecological diversity that are directly derived from them. In contrast, phylogeny-based metrics may offer greater consistency; here, we argue and demonstrate that they provide a more robust framework for eDNA-based biodiversity assessment.

## Why Phylogenetic Diversity Matters in eDNA Metabarcoding Studies

Phylogenetic diversity (PD) is a long-established concept, initially formalized by Faith ^23^, representing a phylogeny-based measure of biodiversity. Before 1992, the term was used more descriptively, but Faith’s work transformed it into a quantitative tool for conservation biology. PD relates to assessments of biodiversity interpreted as the variety of life and has been framed as a measure of “option value”, capturing how evolutionary history contributes to future possibilities for responding to environmental change ^24^.

PD and molecular data go well together. Faith and Baker ^25^ demonstrated how COI-based phylogenies could be used to quantify macroinvertebrate diversity in freshwater ecosystems, arguing that DNA barcoding alleviates key limitations of traditional PD approaches, including uncertain species identification, cryptic diversity, and incomplete taxonomic knowledge, while also enabling the identification of regions of high evolutionary endemism ^26^.

This consistency between molecular data and PD assessment extends naturally to eDNA metabarcoding.

By integrating both taxon occurrence and evolutionary relatedness, PD remains informative across varying levels of taxonomic resolution, enabling robust analyses even when eDNA-based assignments do not reach the species level. While, on one hand, taxonomic inventories and their associated diversity metrics may vary substantially due to taxon non-detection or incomplete species-level identification, and, on the other hand, functional diversity is constrained by the unknown life stage of detected organisms and by strong dependence on selected life-history traits, PD estimates are less sensitive to these limitations.

Moreover, PD computations capitalize on the core strengths of eDNA metabarcoding, notably its potentially broad taxonomic coverage and integrative capacity ^27^, as well as its ability to detect rare, elusive, or cryptic taxa that are frequently overlooked by conventional survey methods ^3,28,29^.

Explicitly incorporating PD into eDNA metabarcoding analyses therefore seems to be particularly relevant, both to reduce experimental variability and to describe biodiversity patterns, helping also to decipher the evolutionary and ecological processes that shape community structures and their capacity to respond to environmental change. Although it represents a novel way of exploiting eDNA metabarcoding data, phylogenetic diversity (PD) remains only sporadically applied in vertebrate-focused studies (**Appendix 1**), while it is far more commonly used in community-level and microbial ecology. To date, only a few eDNA metabarcoding studies targeting aquatic vertebrates have explored PD-based approaches to characterize community structure and evolutionary patterns. All of these studies focus on fish, two of them include elasmobranchs, and only two address vertebrates as a whole^30,31^.

PD metrics inform on a key dimension of biodiversity and are particularly informative for detecting non-random patterns of community structure, resulting from the accumulation of closely related taxa (phylogenetic clustering) or, conversely, from the co-occurrence of distantly related lineages (phylogenetic overdispersion). These patterns are directly relevant for eDNA-based ecosystem monitoring, as they provide insights into processes such as environmental filtering, disturbance, and ecological resilience inferred from detected communities.

### Practical Recipes for Exploring Broad Phylogenetic Patterns Using eDNA Metabarcoding

In practice, PD can be estimated either by anchoring detected taxa to existing phylogenetic backbones or through taxonomy-free approaches that reconstruct trees directly from molecular units such as (M)OTUs/ASVs ^28^. DNA-based bioassessment has explored a range of taxonomy-free strategies, including OTU-based indices, supervised machine-learning pipelines applied to genetic data ^32,33^, and phylogenetic inference of ecological profiles from sequence clusters ^34^. These approaches have proven particularly effective for microorganisms and micro-eukaryotes ^35,36^. However, for macro-organisms, and especially vertebrates, taxonomy-free approaches remain less effective, particularly in applied conservation contexts where species continue to dominate management frameworks and policy discourse. As emphasized by Maclaurin and Sterelny ^37^, species remain the “primary currency of conservation,” despite conceptual ambiguities, taxonomic instability, and limited theoretical justification. In this context, PD offers a powerful compromise: it retains a strong evolutionary foundation while remaining compatible with species-based conservation paradigms.

A central challenge in exploiting the results of eDNA metabarcoding is determining which measures of PD are the most informative, robust and applicable. The so-called “jungle of indices*”* ^38^ encompasses at least 70 different phylogenetic diversity metrics, reflecting both the proliferation of PD measures and ongoing confusion about their mathematical and ecological interpretations ^39^. A unifying framework groups these metrics into three fundamental dimensions ^40^: richness (the sum of accumulated phylogenetic differences), divergence (the mean relatedness among taxa), and regularity (the variance in relatedness across taxa).

In most eDNA studies (**Appendix 1**), the metric of choice is Faith’s PD ^23^, a richness-based metric that quantifies the amount of evolutionary history represented within a biological community ^28^. It is computed as the sum of branch lengths in the phylogeny of observed taxa ^9,41–43^. Empirical evidence suggests that ecological assemblages with higher Faith’s PD tend to exhibit greater productivity and enhanced stability over time ^44^. However, Faith’s PD is a concave function ^45^, tightly correlated with taxonomic richness: adding taxa can increase Faith’s PD but never decrease it, and it rises monotonically with sampling effort.

Beyond richness-based approaches, metrics of phylogenetic divergence have been also widely used in community ecology, particularly the Mean Pairwise Distance (MPD) and the Mean Nearest Taxon Distance (MNTD) ^46–48^. MPD is generally considered more sensitive to tree-wide patterns of phylogenetic clustering or overdispersion, reflecting overall relatedness among co-occurring taxa, whereas MNTD is more sensitive to tip-level structure, capturing clustering or evenness among closely related species near the leaves of the phylogeny ^49^. Although these approaches are conceptually related to Faith’s PD, they shift the focus toward phylogenetic divergence, which is of particular interest because more even spacing of taxa in phylogenetic space is associated with greater functional trait differences and has been linked to enhanced niche complementarity and ecosystem functioning ^44,50,51^.

Complementing divergence metrics, phylogenetic regularity can be quantified using the Variance in Pairwise Distances (VPD), which measures the variability of phylogenetic distances among taxa within a community ^52^. Low variance indicates a more regular and even distribution of lineages, whereas high variance reflects uneven spacing or phylogenetic clustering ^53^, potentially signaling different community assembly processes and levels of functional redundancy.

To assess the significance of these metrics, a few studies (**Appendix 1**) computed standardized effect sizes (SES), which compare observed Faith’s PD, MPD, MNTD or VPD values to null expectations. SES are typically computed from null distributions generated by repeated randomizations (often 1,000), in which species identities are shuffled across the tips of a pruned phylogenetic tree (derived from the original tree and restricted to the set of taxa detected across all samples in the dataset) while preserving community richness. This approach allows departures from random phylogenetic structure to be quantified in a comparable way across communities ^47,54^.

Albeit highly relevant, SES-based patterns can be spatially heterogeneous and difficult to interpret ^55^, particularly in applied or conservation-oriented contexts, where translating deviations from null expectations into management-relevant insights remains challenging ^56^.

To overcome the strong influence of taxonomic richness on phylogenetic metrics, Nipperess and Matsen ^57^ proposed to calculate Faith’s PD accumulation curves using an exact analytical formula. This approach estimates the expected mean and variance of Faith’s PD at each step of the accumulation curve, allowing the calculation of standardized Faith’s PD values at given levels of taxonomic richness, thereby reducing the confounding effect of richness on phylogenetic diversity estimates. Building on this idea, Nipperess ^45^ introduced ΔPD, defined as the difference between Faith’s PD at m = 2 and Faith’s PD at m = 1 (i.e., ΔPD = PD_2_ ™ PD_1_), representing the expected gain in Faith’s PD between the first and second accumulation units, with each accumulation unit corresponding to one randomly drawn taxon. This metric of phylogenetic divergence is intended to capture how evenly taxa are spread across the phylogeny ^45^.

### Decoding eDNA Metabarcoding Data with Phylogenetic Diversity Metrics: Case studies Across Marine Ecosystems

Between 2021 and 2025, we have conducted five campaigns to inventory marine vertebrate biodiversity using eDNA (**Appendix 2**). These campaigns took place in diverse coastal environments (tropical, polar, and temperate) yet sharing a common conceptual framework centered on the study of Urbanized Marine Ecosystems (UME; ^58–60^). They all relied on direct filtering of sea water during 30 min transects, with two samples collected per transect (port and starboard), followed by a metabarcoding process and careful taxonomic assignation ^22^. All results have been published as freely accessible datasets (**Appendix 2**).

We analyzed these data sets in parallel using the same computational protocol, designed to calculate a complementary panel of phylogenetic metrics (code is available on GitHub: Xhttps://github.com/HaderleRachel/A-phylogenetic-turn-for-eDNA-metabarcoding): Faith’s PD, MPD, MNTD, VPD, their SES values, as well as ΔPD.

A comprehensive reference phylogeny for vertebrates was constructed using *U*.*PhyloMaker* ^61^, integrating megatrees for teleosts ^62^, birds ^63^, and mammals ^64^. Faith’s PD accumulation curves were computed with the *PDcalc* package, and ΔPD was extracted as PD_2_ ™ PD_1_.

Taxonomic richness varied strongly among sites (**Figure 1**), with the highest values at GPMG_Har_core and other highly anthropized sites (Abu Dhabi), and the lowest in Svalbard and at the leeward coast of Guadeloupe (LwC). While this partly reflects the classical latitudinal biodiversity gradient, it also suggests higher apparent diversity in anthropized environments (UME). Faith’s PD closely followed taxonomic richness. In contrast, divergence and regularity metrics (MPD, MNTD, ΔPD, VPD) showed the opposite trend, with highest values in Svalbard, whereas Grand Port Maritime de la Guadeloupe (GPMG) and Abu Dhabi sites combined high richness and Faith’s PD with low divergence, reflecting strong phylogenetic clustering.

**Figure 1:**
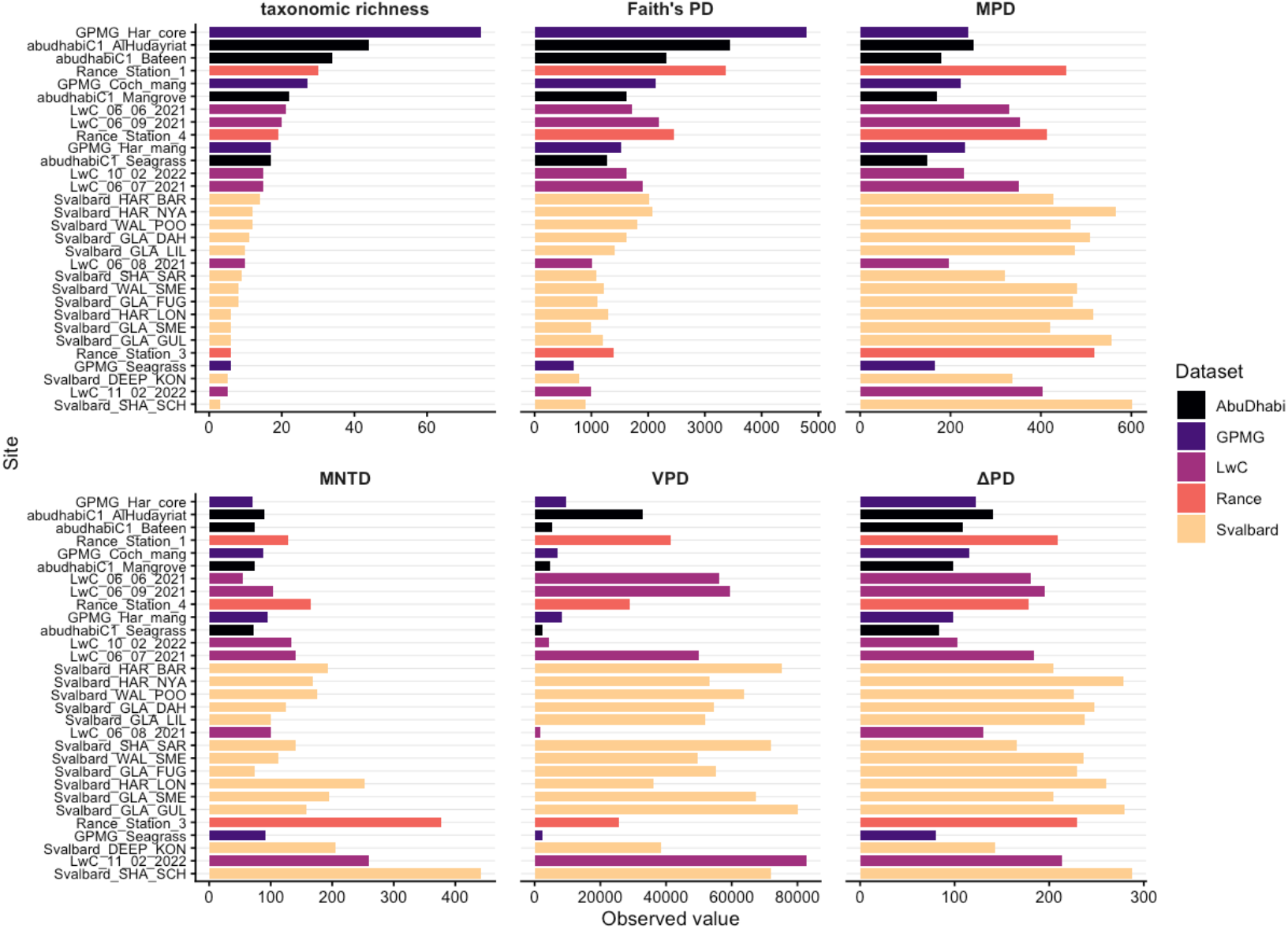
Observed values of taxonomic richness, Faith’s PD, MPD, MNTD, VPD, and ΔPD calculated for each selected site, with colors indicating the corresponding dataset.

SES values (**Figure 2**) confirmed these patterns. GPMG and Abu Dhabi sites consistently showed significantly lower-than-expected PD across metrics. In contrast, Svalbard and Rance sites exhibited positive SES values, consistent with phylogenetic overdispersion. LwC sites were generally close to null expectations.

**Figure 2:**
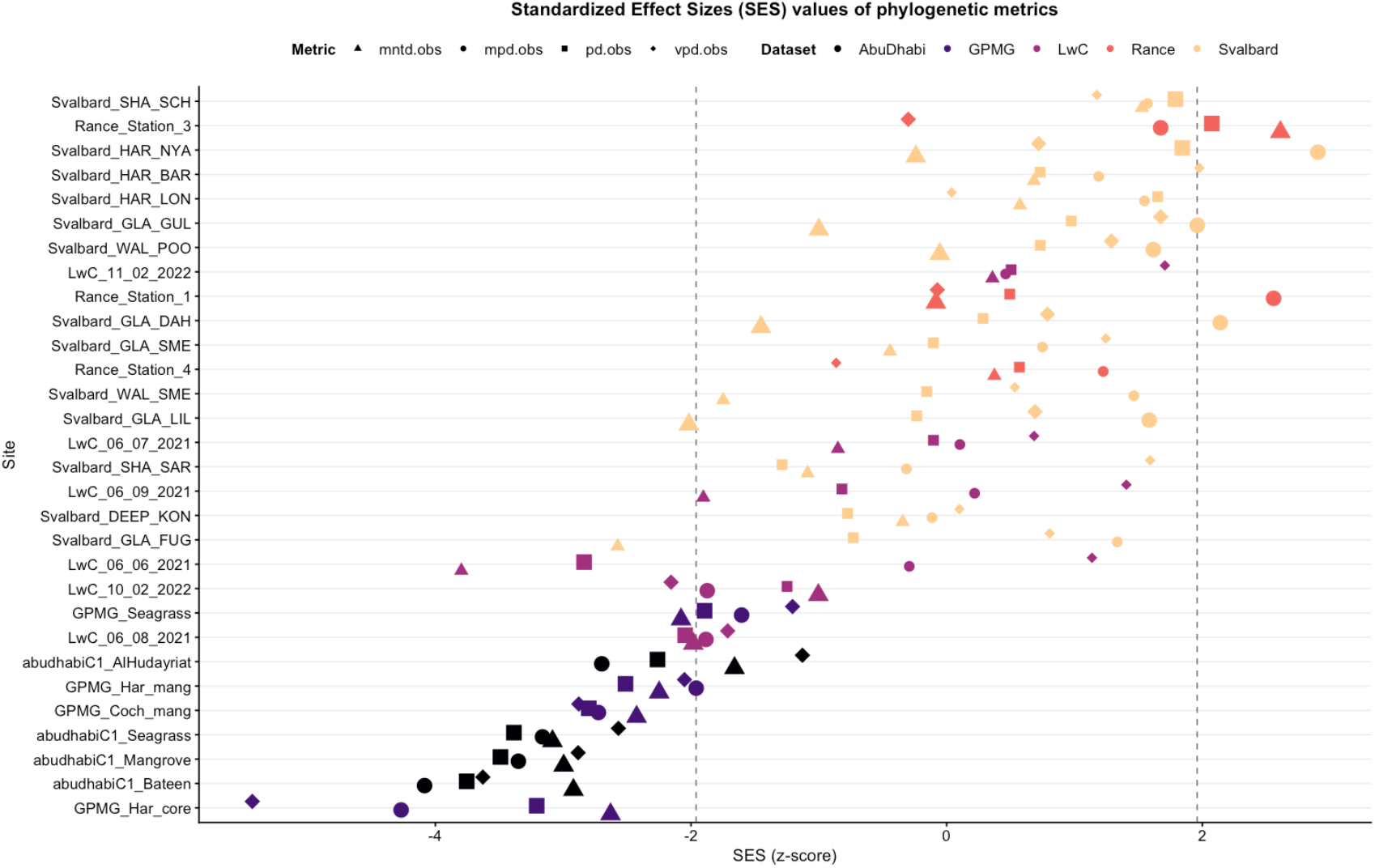
Standardized effect sizes (SES) of phylogenetic diversity metrics (Faith’s PD, MPD, MNTD, and VPD) across sites, with colors representing datasets, shapes indicating each metric, and larger points highlighting values significantly different from expectations based on taxonomic richness.

When sites were pooled by anthropization (**Figure 3**), the “UME” category harbored more taxa and higher Faith’s PD, while the “non-UME” category showed higher divergence and variance metrics. SES values were consistently lower than expected in UME sites and higher in non-UME sites, indicating that anthropized environments concentrate closely related taxa within a narrower portion of evolutionary history. Detailed results are in **Appendix 3**.

**Figure 3:**
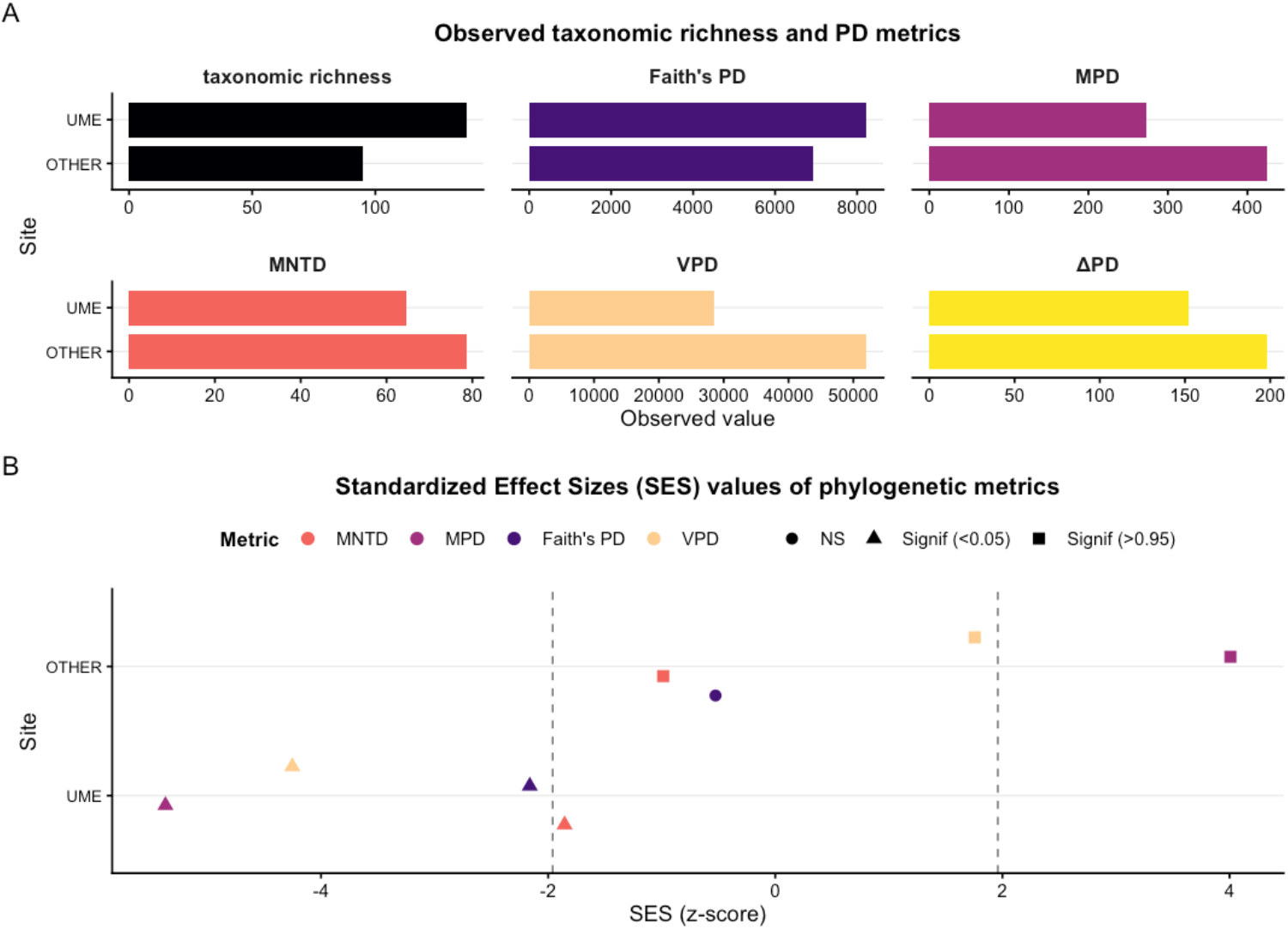
Observed values of taxonomic richness, Faith’s PD, MPD, MNTD, VPD, and ΔPD for the two site groupings (“UME” and “Other”), together with the corresponding standardized effect sizes (SES) for PD metrics.

### A Phylogenetic Turn for eDNA Metabarcoding

eDNA metabarcoding has rapidly become a cornerstone of marine biodiversity monitoring, offering unprecedented taxonomic monitoring coverage across ecosystems. Yet, to fully realize its potential, we argue that a phylogenetic turn is needed.

Moving beyond species lists and richness counts, PD metrics provide a more robust and biologically meaningful view of biodiversity, capturing the evolutionary structure of communities in ways that are less sensitive to common eDNA metabarcoding biases while leveraging the intrinsic strengths of molecular data ^23,24,28^. By quantifying evolutionary history and the “option value” of biodiversity, they highlight how communities may respond to future environmental change. They obviously outperform metrics of functional diversity, when functional roles are poorly known or strongly affected by the blind spots of eDNA metabarcoding, PD is a robust conservation surrogate ^23,65^. Together, these properties make PD metrics highly informative for understanding biodiversity and human impacts, including community assembly, environmental filtering, and ecosystem resilience.

In restored habitats, such as seagrass and mangrove sites in the Grand Port Maritime de la Guadeloupe, PD metrics reveal clear ecological signals: despite apparent recovery, these sites show strong phylogenetic clustering and significantly lower-than-expected PD metrics, consistent with colonization by closely related opportunistic taxa, particularly cryptobenthic teleosts ^9,66^. This sensitivity extends across ecosystems, from polar to tropical environments: species-poor Svalbard sites display phylogenetically overdispersed assemblages, whereas UME concentrate many taxa within narrow evolutionary lineages, revealing consistent patterns of clustering versus divergence and exposing anthropogenic impacts that richness alone would miss ^44,50^.

However, PD remains, at this stage, a set of metrics rather than operational indicators. While highly effective for scientific assessments of biodiversity and human impacts, their translation into decision-support tools requires explicit reference baselines ^67–69^. A central challenge is therefore to turn eDNA results into standardized, goal-oriented indicators for policy and management, anchored to reference states and able to track ecological conditions and trends ^70^. Despite extensive academic uptake (e.g. >4,500 citations of Faith, ^23,55^), PD remains weakly embedded in conservation practice: the EDGE program (^71^; IUCN Resolution WCC-2012-Res-019-EN) remains one of the main applied examples and focuses primarily on species-level distinctiveness rather than community-level evolutionary history. Encouragingly, PD is now entering global policy arenas, including the IPBES Global Assessment ^72^ and the draft Global Biodiversity Framework (CBD, 2022), signaling a growing recognition of its relevance.

Embracing this phylogenetic turn strengthens robustness, ecological relevance, and comparability, and is a necessary step to move from molecular data to policy-relevant biodiversity assessment and fully realize the potential of eDNA metabarcoding.

## Notes

**Conflict of interest disclosure** The authors declare no conflict of interest.

https://github.com/HaderleRachel/A-phylogenetic-turn-for-eDNA-metabarcoding

https://obis.org/dataset/2b47cfee-0233-4bd8-8f12-58a2fc3c5556

https://www.gbif.org/dataset/787386a7-aa69-4f99-ba95-9fd5a700705c

https://www.gbif.org/dataset/fb6939d1-0b75-497e-a54b-2a38a907bf05

https://www.gbif.org/dataset/8c8f29d8-870b-44fd-82c0-c767734e0f27

https://www.gbif.org/dataset/70a53441-4e44-4216-aac3-4e9d02e33212

### Competing Interest Statement

The authors have declared no competing interest.

