## Appendix 1 for "The old pipe gives the sweetest smoke: A phylogenetic turn for eDNA metabarcoding"

Supplementary Material 1: State-of-the-art overview of published studies applying environmental DNA (eDNA) metabarcoding to aquatic vertebrates and explicitly incorporating phylogenetic diversity (PD) metrics.

| **DOI** | **Source** | **Country** | **Marine/Freshwater/Estuary** | **Filtration** | **Targeted taxa** | **Primers** | **PD metrics** | **SES?** | **PD beta-diversity** |
| --- | --- | --- | --- | --- | --- | --- | --- | --- | --- |
| <https://doi.org/10.1371/journal.pwat.0000099> | Manandhar et al., 2023 | Nepal | Freshwater | Indirect | fish | MiFish | Faith's PD | - | - |
| <https://doi.org/10.1038/s41598-024-80907-z> | Deng et al., 2024 | China | Freshwater | Indirect | fish | MiFish | Faith's PD | - | - |
| <https://doi.org/10.1002/ece3.72215> | Li et al., 2025 | China | Freshwater | Indirect | fish | MiFish | Faith's PD | - |  |
| <https://doi.org/10.3390/fishes10090430> | Xu et al., 2025 | China | Freshwater | Indirect | fish | Fish-F4-R4 | Faith's PD, phylogenetic difference index (Delta), weighted phylogenetic difference index (Delta*), phylogenetic homogeneity index (Lambda+) and phylogenetic dispersion index (Delta+) | - | - |
| <https://doi.org/10.1016/j.envres.2025.122661> | Zhang et al., 2025 | China | Freshwater | Indirect | fish | Tele02 | Faith's PD, MPD (NRI), MNTD (NTI) | yes | yes |
| <https://doi.org/10.1101/2025.09.24.676958> | Zhang et al., 2025 | worldwide | Freshwater | Different methods | fish | Different primers | MNTD | - | - |
| <https://doi.org/10.1080/02705060.2025.2541689> | Nneji et al., 2025 | Nigeria | Freshwater/estuary | Indirect | fish | MiFish | Faith's PD | - | - |
| <https://doi.org/10.1002/edn3.140> | Polanco-Fernandez et al., 2020 | Colombia | Marine | Direct | vertebrates (+ focus fish and elasmo) | Vert01, Tele01 and Chon01 | D-stat (phylo signal) to compare with UVC | - | - |
| <https://doi.org/10.1111/cobi.13802> | Marques et al., 2021 | Colombia | Marine | Direct | fish, elasmo | Tele01, Chon01, Vert01 | Faith's PD | - | - |
| <https://doi.org/10.1111/1365-2664.14276> | Dalongeville et al., 2022 | France | Marine | Direct | fish | Tele01 | Faith's PD | - | - |
| <https://doi.org/10.1002/ece3.9212> | Polanco F. et al., 2022 | Curaçao | Marine | Direct | fish | Tele01 | Faith's PD, MPD, MNTD, VPD et VNTD, D-stat (phylo signal) to compare with UVC |  | yes |
| <https://doi.org/10.1002/edn3.305> | Rozansky et al., 2022 | France | Marine | Direct | fish | Tele01 | Faith's PD, MPD, VPD | yes | yes |
| <https://doi.org/10.1111/geb.13698> | Mathon et al., 2023 | worldwide | Marine | Different methods | fish | Tele01 | parwise PD distance, α- and β-diversity Hill number | - | yes |
| <https://doi.org/10.1093/icesjms/fsad139> | Veron et al., 2023 | France | Marine | Direct | fish | Tele01 | Faith's PD, MPD, VPD | yes | yes |
| <https://doi.org/10.1002/edn3.70048> | Chung et al., 2024 | China | Marine | Indirect | fish | Tele02 | Faith's PD, D-stat (phylo signal) to compare with UVC | - | - |
| <https://doi.org/10.1016/j.ecolind.2024.111893> | Jiang et al., 2024 | China | Marine | Indirect | fish | MiFish | Faith's PD | - | yes |
| <https://doi.org/10.3390/fishes9110435> | Wang et al., 2024 | Yellow and Bohai Seas | Marine | Indirect | fish | MiFish | Faith's PD | - | - |
| <https://doi.org/10.1016/j.ecolind.2024.112389> | Zhao et al., 2024 | China | Marine | Indirect | fish | 12S_V5 | Faith's PD | - | - |
| <https://doi.org/10.1002/edn3.70142> | Madon et al., 2025 | Corsica | Marine | Indirect | fish | Tele01 | Faith's PD | - | - |
| <https://doi.org/10.1016/j.marenvres.2025.107439> | Yan et al., 2025 | northwestern Pacific | Marine | Indirect | fish | MiFish | Faith's PD, MPD (NRI), MNTD (NTI) | - | yes |
| <https://doi.org/10.3390/ani15091283> | Zhang et al., 2025 | China | Marine | Indirect | fish | MiFish | Faith's PD, MPD, VPD | yes | - |
| <https://doi.org/10.1002/edn3.232> | Polaco-F. et al., 2021 | France and French Guiana | Marine/estuary | Direct | fish | MiFish and Tele01 | Faith's PD | yes | - |
| <https://doi.org/10.64898/2025.12.01.691502> | Haderlé et al., 2025 | France | Marine, Estuary | Direct | vertebrates | Vert01 | Faith's PD | yes | yes |
