## Appendix 2 for "The old pipe gives the sweetest smoke: A phylogenetic turn for eDNA metabarcoding"

Supplementary Material 2: Overview of the eDNA datasets used in this study, including sampling design, primer sets, sequencing depth, and local anthropogenic pressure level across sites.

| Dataset | Samples | Primers used | Seq. depth | Local anthropization level |
| --- | --- | --- | --- | --- |
| Leeward coast of Guadeloupe (LwC)  (<https://obis.org/dataset/2b47cfee-0233-4bd8-8f12-58a2fc3c5556>) | 4 successive temporal replicates in 2021 and 2 in 2022 | Vert01 ^6^ | 1 mio. reads/sample | Low  (Inside a MPA, approximately 7km from the coast) |
| Rance estuary (<https://www.gbif.org/dataset/787386a7-aa69-4f99-ba95-9fd5a700705c>) | 1 marine site at the mouth of the estuary and 3 estuarine sites | Vert01 ^6^ | 1 mio. reads/sample | Intermediate  (partially protected with anthropogenic pressure) |
| Grand Port Maritime de la Guadeloupe (GPMG)  (<https://www.gbif.org/dataset/fb6939d1-0b75-497e-a54b-2a38a907bf05>) | 1 site at the core of the seaport, 1 located near a degraded mangrove, 2 sites located near restored habitats (mangrove and seagrass) | Vert01 ^6^ | 1 mio. reads/sample | High  (large-scale industrial and commercial port area) |
| Svalbard  (<https://www.gbif.org/dataset/8c8f29d8-870b-44fd-82c0-c767734e0f27>) | 3 seaports, 2 shallow sites, 2 deep sites, 3 walrus haul-out areas, 6 glacier sites | Vert16S ^6^ | 100 000 reads/sample | Low  (protected and sparsely populated Arctic region) |
| Abu Dhabi   (<https://www.gbif.org/dataset/70a53441-4e44-4216-aac3-4e9d02e33212>) | 1 seaport, 1 mangrove, 1 seagrass and 1 channel | Vert01 ^6^ | 1 mio. reads/sample | High  (extensively developed urban area) |
