## Appendix 3 for "The old pipe gives the sweetest smoke: A phylogenetic turn for eDNA metabarcoding"

Supplementary Material 3: Details of the Description and Interpretation of Figures 1, 2 and 3.

Taxonomic richness showed strong contrasts among sites (**Figure 1**). The highest value was observed at GPMG_Har_core (core of the Port of Guadeloupe), followed by the most anthropized Abu Dhabi sites (abudhabiC1_AlHudayriat and abudhabiC1_Bateen), whereas the lowest values were mostly found in Svalbard and at the leeward coast of Guadeloupe (LwC_11/02/2022). Taken alone, this pattern suggests, in addition to the known decreasing pattern of biodiversity from tropical to polar ecosystems ^73,74^, that a higher biodiversity would exist in anthropized sites, with UME appearing richer than likely less impacted locations.

Faith’s PD closely tracked taxonomic richness, as expected from its dependence on species counts (**Figure 1**). In contrast, divergence metrics revealed an opposite pattern. MPD, MNTD and ΔPD were highest in Svalbard sites, and particularly at Svalbard_SHA_SCH, which had the lowest taxonomic richness. VPD followed a similar trend, with relatively high values also observed in the LwC dataset sites. These results indicate that, for example at Svalbard, although species richness was comparatively low, the detected communities spanned a broader portion of the phylogenetic tree. Conversely, anthropized sites such as GPMG_Har_core combined high taxonomic richness and high Faith’s PD with low divergence (MPD, ΔPD, especially MNTD) and low VPD, reflecting strong phylogenetic clustering, notably at the tips of the tree.

SES values (**Figure 2**) are expected to decouple phylogenetic diversity (PD) from taxonomic richness and to assess deviations from null expectations. All heavily anthropized sites (GPMG and Abu Dhabi datasets) consistently exhibited significant SES values across richness-, divergence-, and regularity-based PD metrics, indicating lower PD values than expected. In contrast, Svalbard and Rance sites (notably Rance_Station_3) displayed higher positive SES values, suggesting phylogenetically overdispersed assemblages.

At GPMG, the restored seagrass and mangrove sites (GPMG_Seagrass and GPMG_Coch_mang) showed phylogenetic clustering comparable to that of the highly degraded mangrove site within the port area (GPMG_Har_Mang; **Figure 2**), suggesting colonization by closely related opportunistic species ^75^.

LwC sites generally exhibited SES values close to null expectations, with each sampling date representing a temporal replicate along the same transect over a steep bathymetric drop in the open ocean. However, 3 of the 6 replicates showed at least one PD metric significantly lower than expected (**Figure 2**).

At Svalbard_SHA_SCH, a shallow site with low taxon detection (1 elasmobranch and 3 teleost species), SES values were significantly higher across all four PD metrics (**Figure 2**). Similarly, Svalbard_HAR_NYA, located near the Ny-Ålesund research station, showed detections of 3 bird species, 1 elasmobranch, 3 mammals, and 7 teleost species, also resulting in significative elevated SES values.

Pooling all highly anthropized sites into a “UME” group (GPMG sites; the two Svalbard ports of Longyearbyen, the capital, and Barentsburg, a coal-mining town; the Abu Dhabi marina; and the channel near the marina) versus intermediate-to-low anthropized sites (LwC sites; other Svalbard sites; Abu Dhabi mangrove and seagrass areas; and Rance estuary Station 1) confirmed this pattern (**Figure 3**).

UME-group sites harbored a greater total number of taxa, resulting in higher Faith’s PD, which is strongly correlated with taxonomic richness (**Figure 3**). In contrast, non-UME sites exhibited higher MPD, MNTD, VPD, and ΔPD. SES values were consistently lower than expected in UME sites across all metrics, whereas non-UME sites showed significantly higher-than-expected values for divergence- and variance-based metrics. These results indicate that UME sites concentrate many taxa but represent a narrower and more clustered portion of evolutionary history.
